# A chromosome-scale genome assembly of European Hazel (*Corylus avellana* L.) reveals targets for crop improvement

**DOI:** 10.1101/817577

**Authors:** Stuart J. Lucas, Kadriye Kahraman, Bihter Avşar, Richard J.A. Buggs, Ipek Bilge

## Abstract

European hazelnut (*Corylus avellana* L.) is a tree crop of economic importance worldwide, but especially to northern Turkey, where the majority of production takes place. Hazelnut production is currently challenged by environmental stresses such as a recent outbreak of severe powdery mildew disease; furthermore, allergy to hazelnuts is an increasing health concern in some regions.

In order to provide a foundation for utilizing the available hazelnut genetic resources for crop improvement, we produced the first fully assembled genome sequence and annotation for a hazelnut species, from *Corylus avellana* cv. ‘Tombul’, one of the most important Turkish varieties. A hybrid sequencing strategy combining short reads, long reads and proximity ligation methods enabled us to resolve heterozygous regions and produce a high-quality 370 Mb assembly that agrees closely with cytogenetic studies and genetic maps of the 11 *C. avellana* chromosomes, and covers 97.8% of the estimated genome size. The genome includes 28,409 high-confidence protein-coding genes, over 20,000 of which were functionally annotated based on homology to known plant proteins. We focused particularly on gene families encoding hazelnut allergens, and the MLO proteins that are an important susceptibility factor for powdery mildew. The complete assembly enabled us to differentiate between members of these families and identify novel homologs that may be important in mildew disease and hazelnut allergy. These findings provide examples of how the genome can be used to guide research and develop effective strategies for crop improvement in *C. avellana*.

## Introduction

The genus *Corylus* describes the Hazels, deciduous trees and large shrubs that are widespread throughout the Northern Hemisphere and grown for their edible nuts, wood and ornamental purposes. The most economically significant species is the European Hazel (*Corylus avellana* L.), the nuts of which are known as hazelnuts, filberts or cobnuts and consumed worldwide both directly and as an ingredient in many food and confectionary products. Hazelnuts prefer a mild, damp climate; production is historically concentrated in the Black Sea region of Turkey, which provided ∼65% of the world’s supply in 2017 (FAO 2017). Other major producers include Italy, Azerbaijan, and the USA, and in recent years several other countries have begun actively developing their hazelnut industry, such as China, Georgia, Iran and Chile.

In spite of its widespread use, genetic improvement of *C. avellana* as a crop has been largely limited to the American Pacific Northwest, where the devastating fungal disease, Eastern Filbert Blight, prompted a successful effort to identify and breed for genetic sources of disease resistance (Molnar and Capik 2012; Sathuvalli et al. 2017). In Turkey and elsewhere, hazelnut production is severely affected by abiotic stresses such as frost or drought, and by emerging phytopathogens such as *Erysiphe corylacearum* (Ustaoğlu 2012; Sezer et al. 2017). Over the last 3-5 years, this powdery mildew fungus has become ubiquitous in orchards in Turkey and Georgia, and controlling the disease requires repeated and costly fungicide spraying. Therefore, sources of genetic resistance to powdery mildew are urgently required. In other crop species including wheat, barley, tomato, pea & grapevine, knockdown/knockout of susceptibility genes belonging to the *Mildew Locus O* (MLO) family has been shown to confer resistance to powdery mildew fungi (Acevedo-Garcia et al. 2014). Identification of paralogous genes in *C. avellana* could suggest a target for developing resistant cultivars.

Moreover, in recent years nut allergy has become a well known health problem for a minority of consumers, leading to great interest in the identification of hazelnut allergens and their genes (Costa et al. 2015). To date 11 different allergens, denoted “Cor a” proteins, have been identified and cloned from *C. avellana.* These proteins have diverse structure and functions and some, such as Cor a 1, are found in multiple isoforms with varying levels of allergenicity (Lüttkopf et al. 2002). Characterization of the genomic loci from which these proteins originate would be an important step to understanding how they are produced *in vivo*, and a foundation for developing novel and sensitive DNA-based detection methods for these allergens.

As with many tree species, *C. avellana* has a long generation time (up to 8 years to reach full productivity) and also displays sporophytic self-incompatability, with genetically similar individuals unable to pollinate each other (Marinoni et al. 2009). These factors make selecting for many important traits by classical breeding approaches extremely difficult. Therefore, genomic data, which allows the identification many genetic loci simultaneously, has huge potential to support and accelerate research and breeding for *C. avellana*.

Accordingly, a draft genome assembly and transcriptome of the American cultivar “Jefferson”, along with re-sequencing data from 7 further cultivars, have been produced and made publicly available (Rowley et al. 2012, 2018). Transcriptome sequences have also been produced for two wild hazelnut species, *C. heterophylla* Fisch. and *C. mandshurica* (Ma et al. 2013; Chen et al. 2014). In cultivated *C. avellana,* a genetic linkage map has also been developed (Mehlenbacher et al. 2006), which has been improved by addition of SSR markers developed from both enrichment libraries and the available genome and transcriptome data (Gürcan et al. 2010; Gürcan and Mehlenbacher 2010; Colburn et al. 2017; Bhattarai and Mehlenbacher 2017). Recently, Genotyping-by-Sequencing has been used to generate thousands of SNP markers for a cross between two European cultivars (Tonda Gentile della Langhe x Merveille di Bollwiller), enabling the first genetic mapping of a quantitative trait - time of leaf budburst - in *C. avellana* (Marinoni et al. 2018). Also using a partial genome sequencing approach, novel SSR markers have been developed and used to characterize genetic diversity between Turkish and European hazelnut varieties (Öztürk et al. 2018).

While the studies mentioned above provide essential resources for identification of genes and molecular markers in hazelnut, there is still a need for a reference quality genome sequence of *C. avellana*, in order to identify structural relationships between genes and facilitate rapid mapping of candidate genes from molecular markers for traits of interest. In this study, using the Turkish cultivar ‘Tombul’, we apply a hybrid next-generation sequencing strategy combining short-read, long-read and physical proximity sequencing to generate a *de novo* chromosome-scale genome assembly consisting of 11 pseudomolecules with a total length of 370 Mb. These pseudomolecules are compared to and found to be highly consistent with previous cytogenetic data and genetic maps of the *C. avellana* genome, indicating that they represent a near-complete genome sequence. We also produce a full annotation of the genome sequence, with a detailed analysis of genes and other functional elements predicted to be involved in disease resistance and the production of hazelnut allergens.

## Results

### A hybrid sequencing approach facilitates complete assembly of the hazelnut genome

For an initial survey of the ‘Tombul’ hazelnut genome, we obtained high-coverage Illumina 150 bp paired-end reads for their low error rate and cost-effectiveness. As previously reported (Rowley et al. 2018), this allowed us to produce a *de novo* draft genome assembly; however, this assembly was highly fragmented and 25-30% larger than previous estimates of the *C. avellana* genome size (378 Mb, calculated from flow cytometry data). This could be explained by the heterozygous regions of the genome being assembled twice into separate contigs; accordingly, Benchmarking Universal Single-Copy Orthologs analysis (BUSCO v3) (Waterhouse et al. 2018) found that 25% of highly conserved single-copy genes from land plants (360/1440) were duplicated.

Therefore, we improved the genome assembly by incorporating low-coverage, long single-molecule reads (Oxford NanoPore), and information about physically adjacent sequences produced using proximity ligation sequencing (Dovetail Genomics). These two approaches were combined with the Illumina data separately and together, in order to assess their relative contributions to the final assembly (Table 1). A hybrid assembly of the Illumina and NanoPore data (Illumina + NP) gave 12,557 scaffolds with a total length of 383.1 Mb, comparable to the expected genome size. Although the NanoPore reads were at relatively low genome coverage (9.3x) and have a high base error rate, they enabled the assembly of scaffolds ∼10-fold larger than Illumina-only across the size distribution (Supplementary Data). The Illumina + NP hybrid assembly also eliminated duplicated sequences and the large majority of gaps in assembled scaffolds, compared to the Illumina-only assembly. However, this assembly was still too fragmented to allow large-scale structural comparisons, for example with hazelnut genetic maps and other genomes from other species. In both genome assemblies, the observed GC content was 36%.

**Table 1.** Genome sequencing and assembly statistics.

| Sequencing technology | No. of Reads | Library design | Total read length (Gb) | Sequence coverage<sup>1*</sup> |
| --- | --- | --- | --- | --- |
| **Illumina PE** | 2 x 136M | Paired end, 700-800 bp insert | 41.1 | 108 x |
| **NanoPore** | 1,351,274 | Single molecule, 1-10 kb reads | 3.53 | 9.3 x |
|  |  |  | Insert size range | Physical coverage<sup>2*</sup> |
| **Dovetail Chicago** | 2 x 221M | Proximity ligated | 1-100 kb | 225 x |
| **Dovetail HiC** | 2 x 242M | Proximity ligated | 10 kb – 10 Mb | 3,447 x |
| **Assembly Statistics** |  |  |  |  |
|  | Illumina only | Illumina + Dovetail | Illumina + NP | Illumina + NP + Dovetail |
| No. of scaffolds (>1kb) | 89,427 | 32,741 | 12,557 | 2,206 |
| Scaffold N50 / L50 | 20927 / 7.32 kb | 12 / 13.33 Mb | 1299 / 78.8 kb | 5 / 36.65 Mb |
| Scaffold N90 / L90 | 149112 / 204 bp | 89585 / 3.4 kb | 5857 / 13.7 kb | 10 / 22.72 Mb |
| Total scaffold size (Mb) | 513.8 | 520.1 | 383.1 | 384.2 |
| % gap bases (N) | 6.01% | 6.45% | 0.18% | 0.47% |
| Largest scaffold | 0.152 Mb | 45.9 Mb | 0.732 Mb | 50.95 Mb |

<sup>1</sup>Average no. of times a genome nucleotide is included in a sequence read
<sup>2</sup>Average no. of times a genome position is included in the region between 2 linked reads
\* Calculated for an estimated genome size of 378 Mbp.

In order to improve the contiguity of the assembly, we carried out proximity ligation sequencing using Dovetail Genomics’ proprietary methods. These generate pairs of linked reads that originate from within the same large DNA fragment (Chicago library) or from physically adjacent nucleosomes in native chromatin (HiC library). The sequences of the linked reads are not assembled directly, but mapped to the scaffolds from a pre-existing genome assembly. The ‘HiRise’ bioinformatic pipeline then uses these links to determine the order and orientation of scaffolds along each pseudomolecule. Adjacent, non-overlapping scaffolds are joined with an arbitrary gap sequence of (N)_100_. As shown in Table 1, incorporation of the Dovetail data enabled pseudomolecules longer than >10 Mb to be assembled both from Illumina-only and Illumina + NP assemblies. However, the large number of small scaffolds in the Illumina + Dovetail assembly remained unassembled, and the duplicate scaffolds were not resolved. In contrast, the Illumina + NP + Dovetail assembly consisted of 11 chromosome-sized pseudomolecules ranging from 22.42 – 50.95 Mb in length (Table 2), in total accounting for 97.8% of the predicted genome size; the remaining unplaced scaffolds were in the size range 1-100 kb. The chromosome-sized pseudomolecules (hereafter ‘chromosomes’ for brevity) were labelled pchr01 – pchr11 in descending order of size. The completeness of the assembly was confirmed by BUSCO analysis; orthologs of 97% of 1440 highly conserved land plant genes were found in the chromosomes (90% complete single copies, 6% complete and duplicated, 1% fragmented). We assessed the large-scale accuracy of the hybrid assembly by comparison with previously published cytogenetic analysis of *C. avellana* chromosomes (Falistocco and Marconi 2013). Falistocco and Marconi confirmed the karyotype of diploid *C. avellana* as 2n=22, and noted that there were 3 distinct size groups of 2 large, 5 medium and 4 small chromosomes. Similarly our hybrid assembly contains 2 chromosomes of ∼50 Mb, 5 ranging from 30-40 Mb, and 4 in the range 22-25 Mb. The aforementioned study also used *in situ* hybridization to locate the 45S & 5S rDNA repeats on one chromosome from the large and small groups respectively; using BLAST, we located a 45S rDNA on pchr02, and the 5S rDNA on pchr11. Furthermore, known *C.avellana* SSR sequences were mapped on to the pseudomolecules and compared with the genetic map of the cross OSU 252.146 x OSU 414.062 (Colburn et al. 2017) (Supplementary Figure 1). Each chromosome only contained SSRs from a single linkage group, meaning that they could be unambiguously identified, and 108/113 (95.6%) of the shared SSRs were co-linear in the 2 datasets, suggesting that there are no large-scale structural differences between the ‘Tombul’ chromosomes and the varieties from which the genetic map was constructed.

**Table 2.**
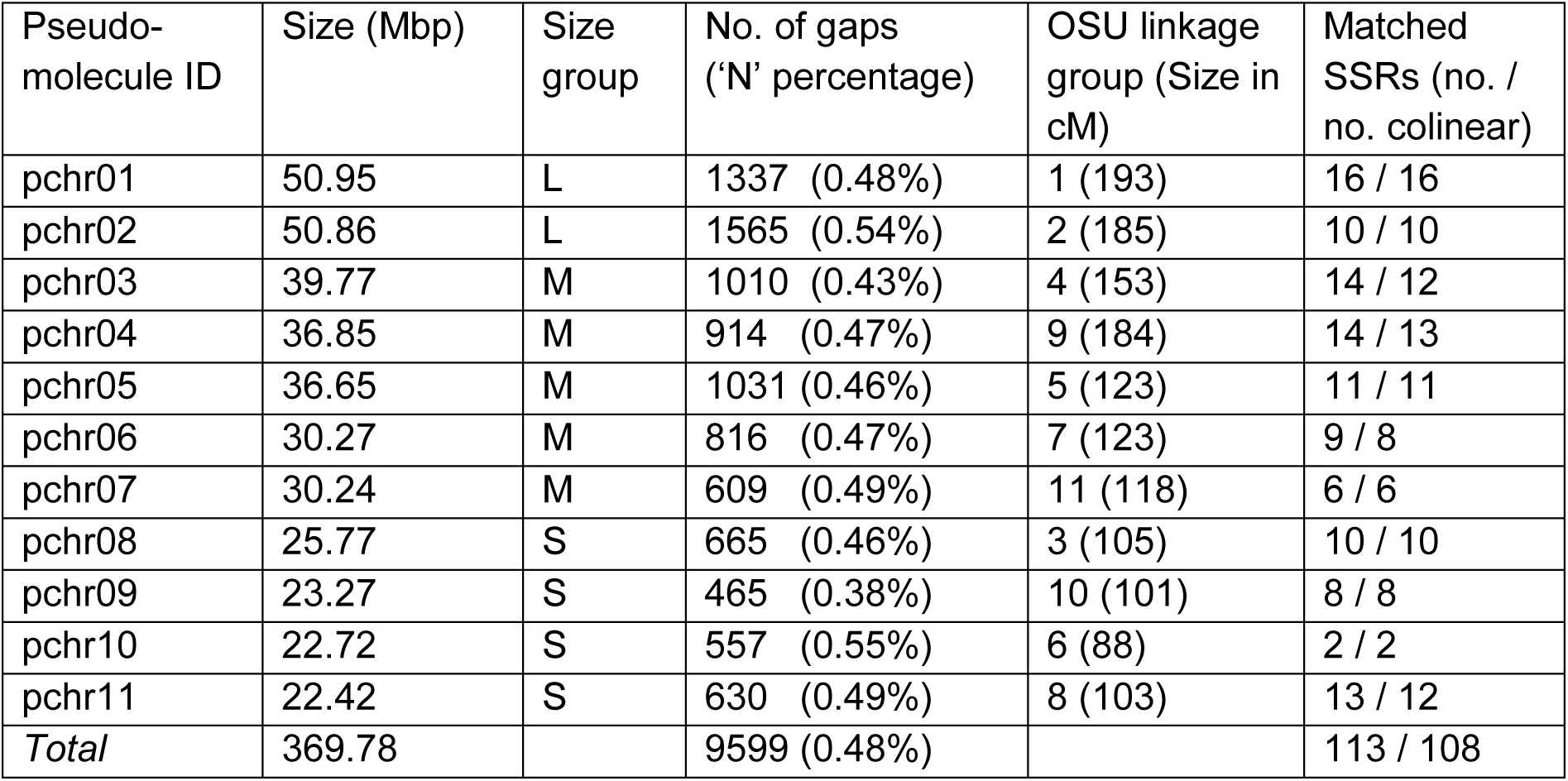
Chromosome pseudomolecule statistics.

| Pseudo-molecule ID | Size (Mbp) | Size group | No. of gaps ('N' percentage) | OSU linkage group (Size in cM) | Matched SSRs (no. / no. colinear) |
| --- | --- | --- | --- | --- | --- |
| pchr01 | 50.95 | L | 1337 (0.48%) | 1 (193) | 16 / 16 |
| pchr02 | 50.86 | L | 1565 (0.54%) | 2 (185) | 10 / 10 |
| pchr03 | 39.77 | M | 1010 (0.43%) | 4 (153) | 14 / 12 |
| pchr04 | 36.85 | M | 914 (0.47%) | 9 (184) | 14 / 13 |
| pchr05 | 36.65 | M | 1031 (0.46%) | 5 (123) | 11 / 11 |
| pchr06 | 30.27 | M | 816 (0.47%) | 7 (123) | 9 / 8 |
| pchr07 | 30.24 | M | 609 (0.49%) | 11 (118) | 6 / 6 |
| pchr08 | 25.77 | S | 665 (0.46%) | 3 (105) | 10 / 10 |
| pchr09 | 23.27 | S | 465 (0.38%) | 10 (101) | 8 / 8 |
| pchr10 | 22.72 | S | 557 (0.55%) | 6 (88) | 2 / 2 |
| pchr11 | 22.42 | S | 630 (0.49%) | 8 (103) | 13 / 12 |
| *Total* | 369.78 |  | 9599 (0.48%) |  | 113 / 108 |

Taken together, these data indicate that the chromosome sequences presented here are consistent with what is known about the physical structure of the hazelnut genome, while the few differences may be the result of local translocations specific to the variety sequenced here. Therefore, we propose that these data can be used as a reference genome for ongoing studies, especially for Turkish hazelnut varieties.

### Functional annotation of the hazelnut genome

The chromosome sequences were annotated as described in detail below to identify repetitive sequences and functional elements such as rRNA, tRNA, miRNA and protein-coding genes (Figure 1). As is typical for eukaryotic genomes, transcribed genes were more abundant towards the ends of the chromosomes, while repetitive DNA content was concentrated near the centromeres. Protein-coding genes were also clustered into orthologous groups using OrthoMCL(Fischer et al. 2011). It was observed that the large number of orthologs (4,324) were found as adjacent copies or clusters, suggesting that these gene families have undergone local tandem duplications. Conserved blocks of 3 or more genes from different orthologous groups were also identified across the genome, and it was noted that most of these repeated blocks were also found in fairly close proximity to each other within a single chromosome, with only a handful showing evidence of possible historical inter-chromosomal duplications (Fig. 1, innermost tracks). The long arm of pchr01, pchr02 and parts of pchr10 seemed to have a higher density of duplicated gene blocks than the rest of the genome, suggesting that these regions may contain recombination hotspots

**Figure 1.**
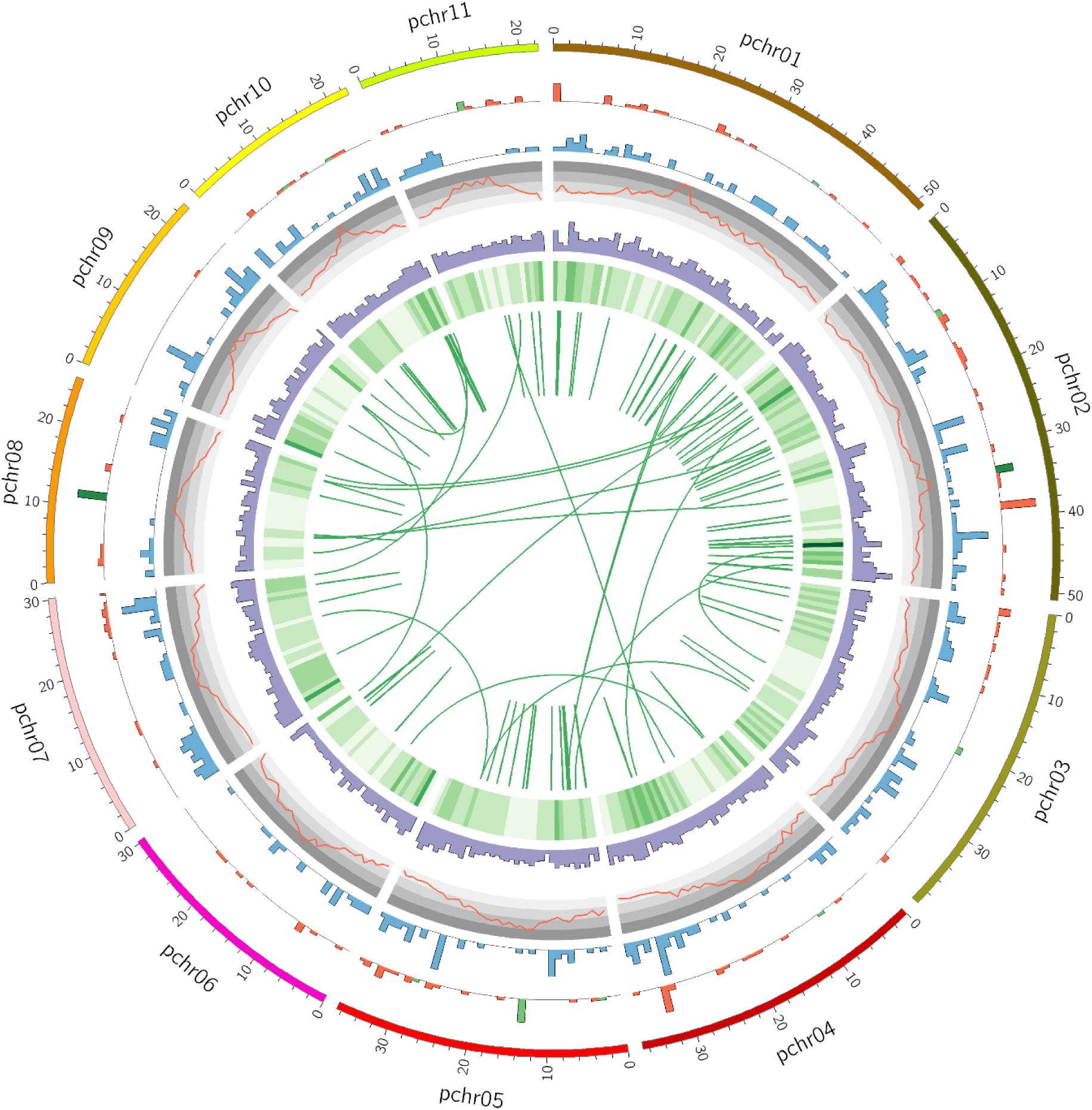
Circular plot of the *C. avellana* cv. Tombul genome summarizing functional features (Detail in Supplementary Tables 2-9). Working in from the outside: i. Ideogram of pseudochromosomes, with lengths marked in Mbp, ii. Histogram of miRNA (red), 5S rRNA (green) and 45S rRNA (dark green) gene density, iii. Histogram of tRNA genes (blue), iv. Line graph of repetitive content as % of total sequence (background shading from light to dark grey indicates the inter-quartile ranges), v. Histogram of protein-coding gene density (purple), vi. Heatmap of tandem gene duplications (darker green indicates more duplications), vii. Links showing repeated blocks of 3 or more adjacent gene paralogs, indicating past translocations and duplications.

### Repetitive landscape of the hazelnut genome

Initial screening of the genome assembly with repetitive elements previously annotated in other eudicots detected few matches (11.87% of the genome), suggesting that the majority of repetitive elements in the genome are lineage-specific. Therefore, prediction tools were used to generate a database of *Corylus*-specific transposable elements based on known structural features of each type (Supplementary Information). When these were included, 35.72% of the entire genome assembly was found to consist of interspersed repeats, while a further 2.41% was made up of simple repeats and low-complexity sequences (Supplementary Tables 1 & 2). Repetitive content varied widely along each chromosome, from 25% or less near the ends to 75-90% in the pericentromeric regions. Over 92.7% of the repetitive DNA comprised retroelements with long terminal repeats (LTRs). Over half of these sequences were incomplete LTR elements, with internal deletions and too much sequence diversification to positively assign them to a repeat family. Of the remainder, Copia elements were almost twice as abundant as Gypsy elements (Supplementary Figure 2). In the *Betula pendula* genome, some classes both of DNA transposons and non-autonomous retroelements were highly abundant (Salojärvi et al. 2017); however, this was not the case in *C. avellana,* suggesting that expansion of these families took place only in the *Betula* lineage.

Taken together, our observations suggest that the repetitive landscape of hazelnut is relatively static, with few elements being highly active in recent evolutionary history.

### Annotation of *C. avellana* functional RNAs

Genes coding for proteins and functional non-coding RNAs were predicted and annotated as described below. A total of 477 predicted tRNA genes and 40 tRNA pseudogenes were distributed across all the chromosomes (Fig. 1, Supplementary Table 3), representing all 20 amino acids and 54 of the possible anticodons. These included ten putative suppressor tRNAs, with anti-codons complementary to the TGA (9) or TAA (1) stop codons. tRNA types and their codon preferences were also analyzed in three other tree species (Supplementary Table 4 & Figure 3). Ribosomal RNA genes were found on pchr02 & pchr08 (45S rDNA) and pchr05 & pchr11 (5S rDNA), while ribosomal proteins were distributed among all chromosomes except pchr11 (Supplementary Tables 5 & 6).

MicroRNAs are ubiquitous post-transcriptional regulators in plants and a previous study identified putative miRNA genes in the draft *C. avellana* cv. Jefferson genome (Avsar and Aliabadi 2017). In the Tombul genome, 153 putative conserved miRNA genes were annotated, including members of 52 different miRNA families (Supplementary Table 7). The majority (95/153) of predicted mature miRNA sequences were 21 nt in length, and the most abundant miRNAs were the well-characterized miR156, miR171 & miR399 families, with 16, 12 & 12 candidates respectively (Supplementary Figure 4). Mapping the predicted pre-miRNAs to assembled transcriptome sequences found evidence for expression of 22/52 miRNA families under normal growth conditions, many of which have also been functionally annotated in other species (Table 3).

**Table 3.** miRNA families expressed in *C. avellana* under normal growth conditions.

| miRNA family<br>(loci in genome) | Targets confirmed in other species | Reported Biological<br>Functions |
| --- | --- | --- |
| miR156/157 (16) | Squamosa-promoter Binding Protein (SBP)<br>box transcription factors | Shoot & leaf<br>development |
| miR160 (2) | Auxin Response Factor (ARF) proteins | Auxin signalling, plant<br>development |
| miR162 (1) | Dicer-Like 1 (DCL1) proteins | miRNA processing |
| miR164 (3) | NAC domain transcription factors | Shoot apical meristem<br>formation, abiotic stress |
| miR165/166 (4) | HD-Zip transcription factors | Meristem & leaf<br>development |
| miR167 (5) | Auxin Response Factor (ARF) proteins | Floral development,<br>stress responses |
| miR169 (8) | CCAAT-box binding / NF-Y transcription<br>factors | Abiotic & biotic stress<br>responses |
| miR170/171 (12) | GRAS-domain or SCARECROW-like<br>transcription factors | Root patterning, light &<br>gibberellin signalling |
| miR172 (4) | APETALA2-like transcription factors | Plant development |
| miR319 (4) | TCP-like transcription factors | Leaf development,<br>abiotic stress response |
| miR393 (1) | F-box proteins & bHLH transcription factors | Root development,<br>auxin signalling |
| miR394 (4) | F-box proteins | Plant development,<br>stress responses |
| miR396 (3) | GRF transcription factors, rhodenase-like<br>proteins, kinesin-like protein B | Growth regulation |
| miR403 (3), miR482 (5), miR529 (1), miR1863 (2),<br>miR5021 (7), miR6288 (1), miR8738 (1), miR9752 (1) |  | *No experimentally*<br>*confirmed targets* |

Other potential targets for the miRNAs identified in this study were predicted by searching for miRNA complementary sequences in *C. avellana* transcriptome sequences. Transcripts with potential miRNA target sites were then annotated with GO terms (Supplementary Table 8; Fig. 2A,B). In the Biological Process domain, the greatest number of GO annotations were related to cellular organization, communication and signal transduction, while in the Molecular Function domain nucleic acid binding, kinase activity, and protein binding were most prevalent. These observations suggest that under normal conditions, *C. avellana* miRNAs are primarily involved in the regulation of cell growth and tissue development. The miRNA complement of *C. avellana* cv. Tombul was also compared with that predicted from the draft cv. Jefferson genome and 5 other tree species (Fig. 2C). While there was a well-conserved group of 26 miRNA families common to all the species examined, there were also miRNA families that were unique to each species. Surprisingly, there were 8 miRNA candidates that were predicted only in Tombul but not Jefferson, and 4 for which the reverse was true. Although there were no transcripts for these miRNAs in our dataset, 2 of them (miR1520 & miR7486) were recently identified by small RNA sequencing from dried hazelnuts (Aquilano et al. 2019); further experiments would be useful to confirm whether the other candidates are functional miRNAs. Also of interest are miR1863, miR8148 and miR8738, which were predicted in both Tombul and Jefferson but none of the other tree species, and were supported by transcriptome data. miR1863 was originally identified in the rice genome but has also been reported to be present in melon (*Curcumis melo*) and Norway spruce (*Picea abies*); this is the first time it has been predicted in the Fagales, suggesting that it may have a lineage-specific function. Both miR8148 and miR8738 were found among small RNAs in dried nuts; it has also been demonstrated that some nut miRNAs can interact with genes from the mammalian immune system, such as miR-156c with the TNF-α receptor (Aquilano et al. 2019). In the light of the known allergenicity of hazelnuts, further investigation of these *Corylus*-specific nut miRNAs would be of great interest to see whether they might have any effects on ingestion.

**Figure 2.**
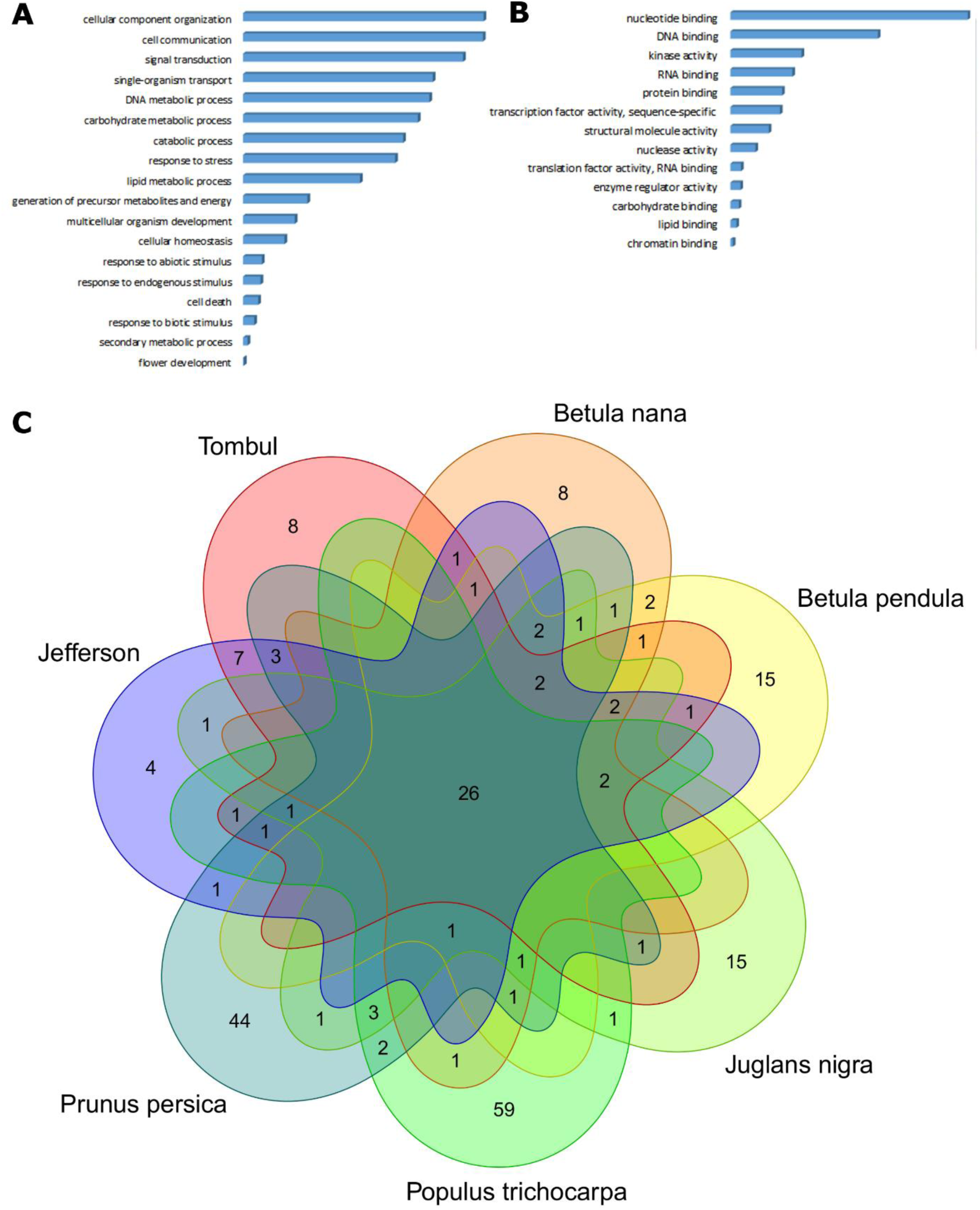
**A, B.** Most abundant GO terms assigned to predicted miRNA target mRNAs in the Biological Process (A) and Molecular Function (B) domains. **C.** Venn diagram of conserved plant miRNA families identified in *C. avellana* cv Tombul and other published tree genomes.

### Gene complement of *C. avellana* cv. Tombul

Using an *ab initio* method, 50,906 gene models were predicted in the repeat-masked pseudochromosomes. These were filtered using Tombul transcriptome sequences to include only transcribed regions, resulting in 28,409 high-confidence protein-coding gene models (Supplementary Table 9). Functional annotation of predicted genes was carried out by sequence similarity to known plant proteins using 3 different strategies; Mercator4 was used to assign predicted protein sequences to MapMan ‘Bins’, while the Trapid web server was used to assign Gene Ontology (GO) terms and search for conserved protein domains. The MapMan Bins represent plant-specific molecular components and pathways; 37.24% of gene models (10,579) were classified by this approach; a further 6,674 were functionally annotated with their closest matching protein, but these had not been assigned to a bin (Supplementary Table 10). The most populous bins were Protein and RNA processing (11.39% & 8.25% of all gene models respectively), followed by Signalling and Stress Responses with 5.36% and 3.98% (Figure 3A). Expansion of specific gene families may indicate functions that have been important in the evolution of *C. avellana.* Therefore, the bin assignments were compared with those for 6 other representative plant species; 62 bins, associated with diverse functions, were identified for which the *C. avellana* genome contains at least 50% more representatives than any of the reference plant genomes (Supplementary Table 11).

**Figure 3.**
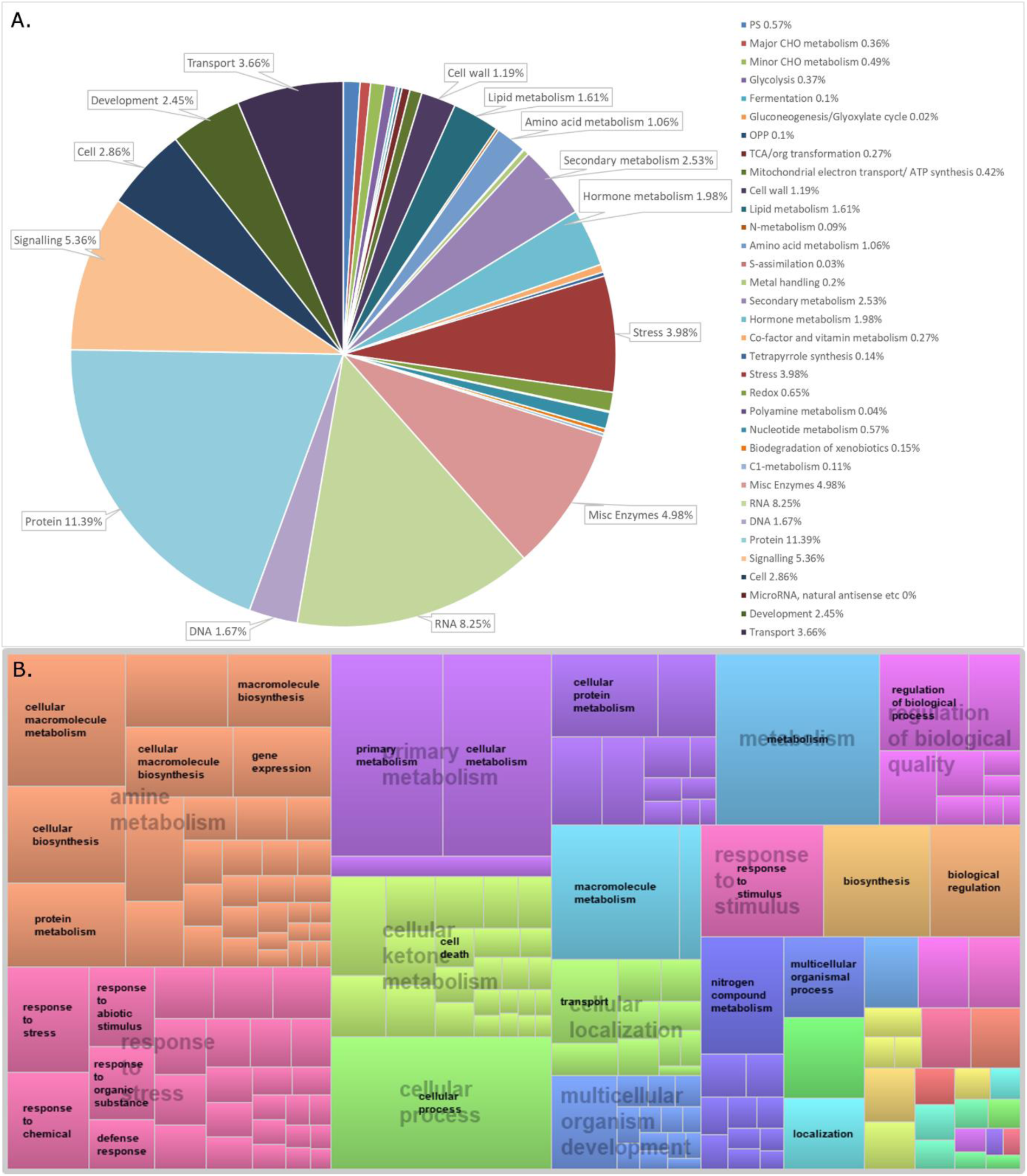
Summary of annotated *C. avellana* gene models, clustered by predicted functions. **A.** Proportion of gene models assigned to the 34 MapMan Bins by Mercator, as % of all high-confidence gene models. Unassigned gene models (42.06%) are not shown. **B.** Treemap view of clustered GO terms in the Biological Process domain. The size of each rectangle is proportional to the number of gene models annotated with this term, and terms clustered in blocks of the same colour on the basis of semantic similarity.

Using the Gene Ontology approach, 69.6% of gene models (20,113) were annotated with one or more GO terms, while 71.8% (20,404) contained at least one conserved protein domain from the InterPro database (Supplementary Tables 9 & 12), discussed more fully in the Supplementary Information. Clustering of similar Biological Process GO terms again found those associated with Protein Metabolism to be most abundant, followed by Stress responses; in addition several other aspects of metabolism, regulation and development were highlighted (Fig.3B). All three annotation approaches identified a large complement of genes encompassing functions that are ubiquitous in plant genomes; however, gene models predicted to be involved in stress responses were notably abundant. Furthermore, almost 8,000 gene models were not annotated by any of these approaches, indicating the need for further study to elucidate their functions.

### The *MLO* gene family in *C. avellana* as a target for powdery mildew resistance

MLO proteins were first identified in barley, where a loss-of-function mutation in the gene Mildew resistance Locus O was found to confer durable resistance to nearly all strains of the barley powdery mildew pathogen, *Blumeria graminis* f.sp. *hordei* (Büschges et al. 1997). Although powdery mildew is caused by different fungal species on each plant host, resistance to mildew infection of *MLO* mutants has since been observed in *A. thaliana*, tomato, and pea, (Acevedo-Garcia et al. 2014) and introduced by gene editing in wheat (Wang et al. 2014). Therefore, this mechanism seems to be functionally conserved across diverse plant species and could present a promising target for developing *E. corylacearum* powdery mildew resistance in hazel.

The annotation pipeline described above identified 24 gene models that had high similarity to the InterPro domain IPR004326, ‘MLO-related protein’. These were examined in detail and manually re-annotated as described in Supplementary Table 13. On the basis of sequence similarity and secondary structure prediction, 11 of these gene models were found to encode full-length MLO proteins, while most of the remainder appeared to be truncated orthologs of the full-length genes. In order to identify those most likely to be involved in powdery mildew infection, the predicted protein sequences were aligned with the original MLO protein from barley (HvMLO), those previously described in *A. thaliana*, and from apple (Pessina et al. 2014), the most closely related species to hazel in which this family has been studied in depth. A phylogenetic tree was used to cluster the sequences, which formed 8 clades, in agreement with previous studies (Figure 4A & Supplementary Figure 5). No *C. avellana* MLO proteins were found in Clade IV, which consists mostly of monocot mildew resistance genes, or Clade VIII, which has only been identified in a subset of Rosaceae species (here represented by MdMLO20). Interestingly, the largest number of *C. avellana MLO* genes (4) fell into Clade V, which also contains all of the dicot *MLO* genes that have been demonstrated to confer susceptibility to powdery mildew infection – of those shown here, *AtMLO2*, *6 & 12*, along with *MdMLO19*. Clade VII was not clearly separated from Clade V in this analysis, suggesting that these genes should also be studied for any potential role in PM.

**Figure 4.**
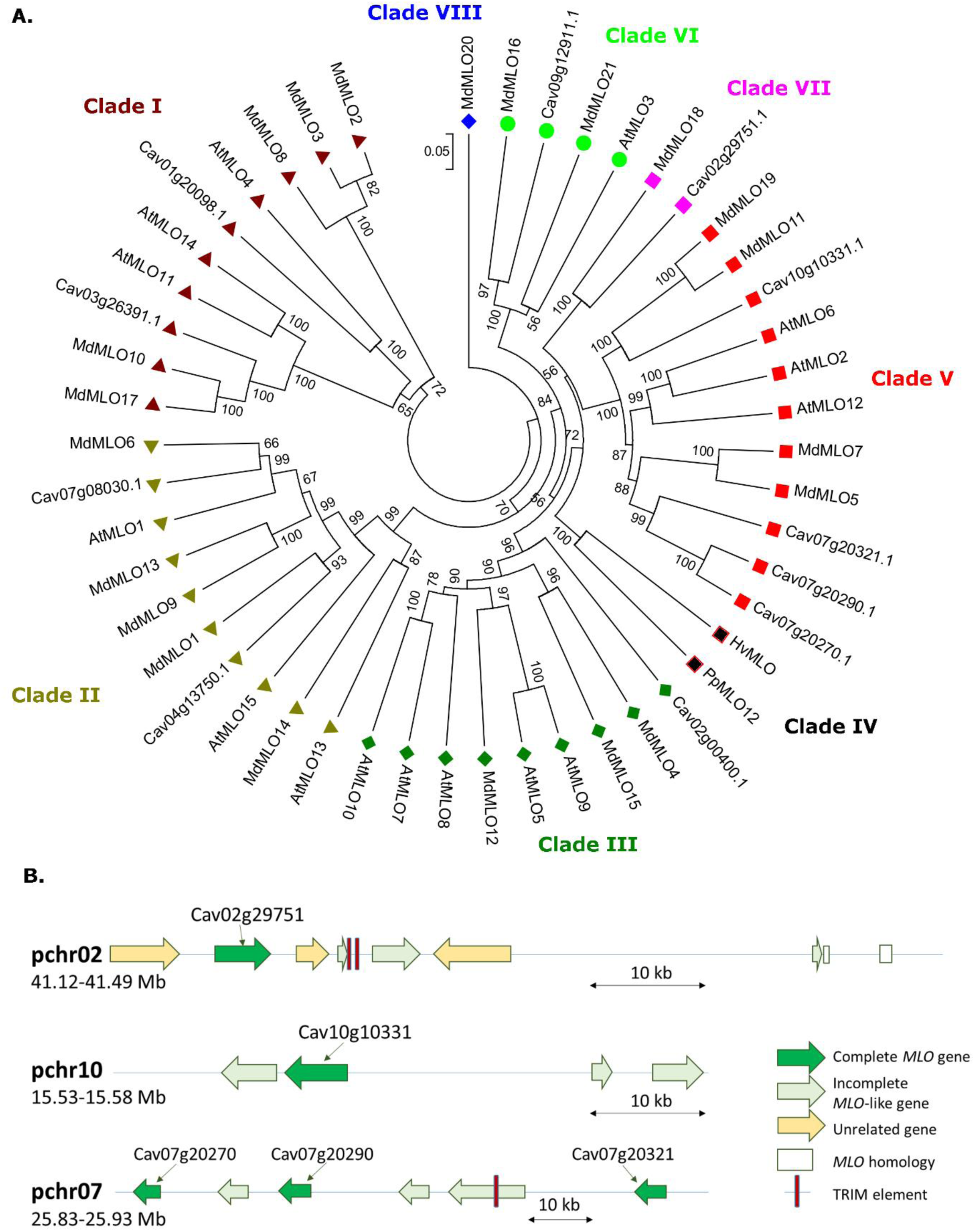
**A.** Phylogenetic clustering of *C. avellana MLO* gene models with those from *A. thaliana* & *Malus domestica*, using the UPGMA approach. Branch lengths are scaled to the no. of amino acid differences per site (p-distance method); node confidence values are % of 1000 bootstrap replications. **B.** Schematic of gene model predictions in pseudochromosome regions containing *MLO* gene models from clades V & VII.

The genome context of the Clade V & VII *MLO* genes was also examined in detail (Figure 4B). Strikingly, all five of these genes had truncated homologs nearby. Two disrupted *MLO* genes with high sequence identity to Cav02g29751 were found in the 50 kb following of the full-length gene. In the first, an insertion of two TRIM elements near the beginning of the gene had split it into two separate open reading frames. In the second, a stop codon truncated the predicted protein after the first 180 amino acids, but the remaining unexpressed exon sequences were still present further downstream. Similarly, Cag10g10331 was found adjacent to a probable tandem duplicate with an N-terminal truncation, while two other truncated *MLO-*like genes, were located within 50 kb on the opposite strand. Finally, the cluster on pchr07 contains three full-length genes that are all closer in homology to each other (Cav07g20270, 20290 & 20231) than any other *MLO* genes, interspersed with multiple truncated gene copies. In one case, two partial MLO-like genes appear to have been spliced together into a single, longer gene, possibly as a result of a TRIM insertion into one of the introns. Taken together, these observations suggest that the Clade V/VII *MLO* genes have undergone repeated tandem duplications followed by degeneration of many of the copies during the development of the hazelnut genome. Most of the other hazelnut *MLO* genes (with the exception of Cav4g13750) did not have degenerate copies in the genome, which may suggest that there has been specific selective pressure for diversification of the Clade V/VII *MLO* genes.

### Genomic insights into hazelnut allergenicity

Hazelnut allergens to date have been identified empirically by screening hazelnut protein extracts with sera from nut allergic patients. Proteins which show specific IgE reactivity were then partially identified by Edman sequencing; this provides enough sequence to design primers and retrieve the complete coding sequence by RT-PCR (Beyer et al. 2002; Schocker et al. 2004). By this approach, to date 11 hazelnut allergens have been identified and recorded in the WHO allergen database (www.allergen.org), while a 12th (Cor a TLP) has been reported but is not yet confirmed (Palacín et al. 2012). We used the published sequences of these allergens to identify their coding genes, and homologs, in our genome annotation (Table 4). We found gene models predicted to encode proteins with 96-100% amino acid identity to all the known allergens; the few variations can be attributed to sequence diversity between hazelnut cultivars.

**Table 4.**
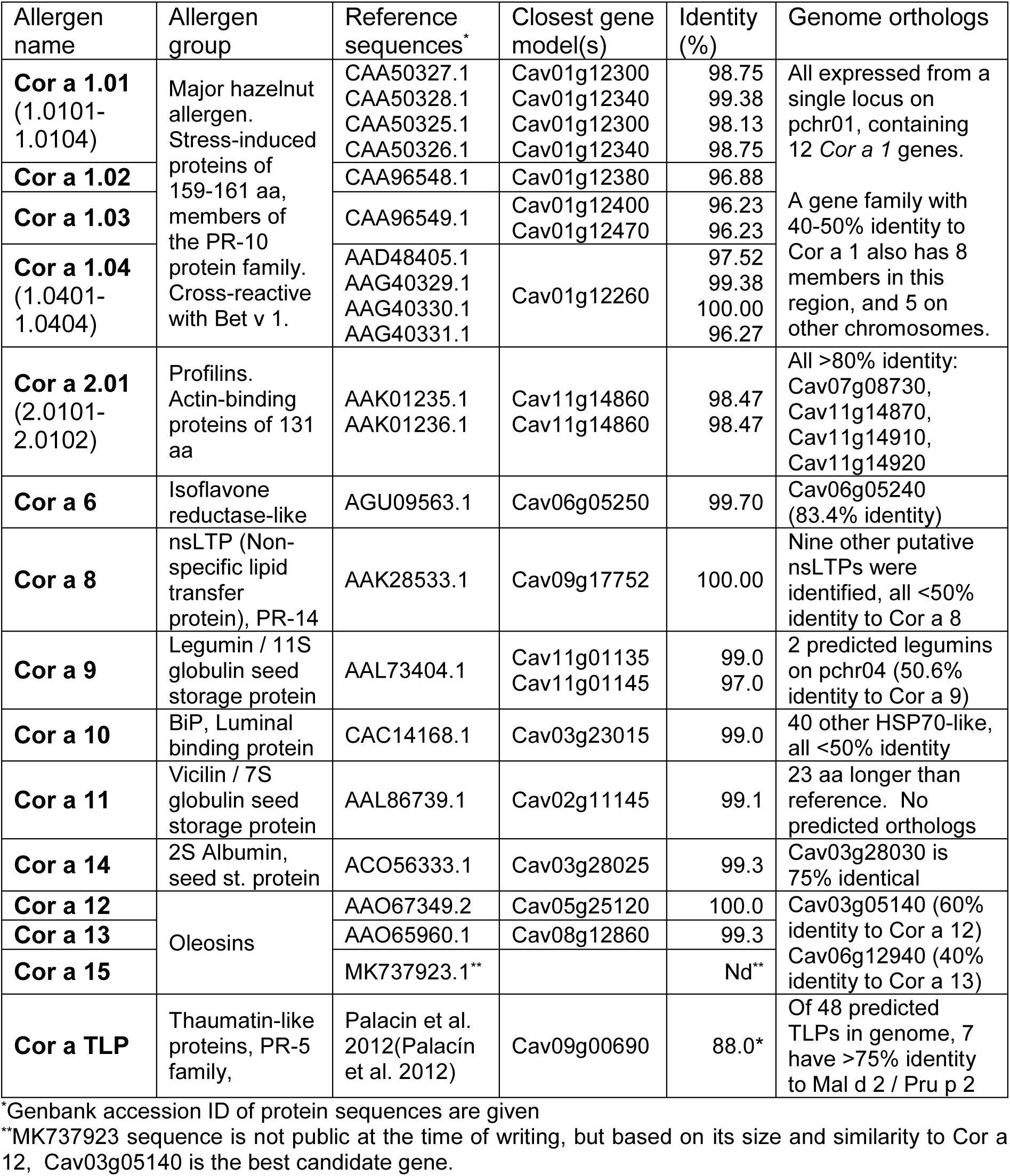
Genes encoding hazelnut allergen proteins in *C. avellana* cv. Tombul.

Food allergens frequently comprise groups of closely related proteins, that show IgE cross-reactivity including across species. ‘Isoallergens’ are defined as proteins from the same species that show cross-reactivity and at least 67% amino acid identity. Within these, ‘isoforms’ are considered to be variants of the same allergen, typically with >90% identity (Chapman et al. 2007). The first and most fully characterized hazelnut allergen is Cor a 1, which is reported to include 4 isoallergens, two of which themselves have 4 isoforms (Breiteneder et al. 1993; Lüttkopf et al. 2002). Cor a 1 is also cross-reactive with paralogs from other Betulaceae species, such as the pollen allergens Bet v 1 from birch, and Car b 1 from hornbeam.

We found that gene models for all the previously reported Cor a 1 proteins were found within a single locus on pchr01 (from 12.3-12.7 Mb); this locus included 20 predicted Cor a 1 homologs, interspersed with unrelated genes. We carried out a phylogenetic comparison of all the predicted Cor a 1 protein sequences (Figure 5), revealing that the relationship between these and the established isoallergen nomenclature is complex. Twelve of the genes were highly conserved with reported Cor a 1 sequences; all of these encoded proteins of 159-161 amino acids in length, and had a fixed 2 exon structure with the intron interrupting codon 62, typical of the Bet v 1 allergens (Hoffmann-Sommergruber et al. 1997). The remaining eight homologs, along with five others found on other chromosomes, formed a group with variable intron-exon structure and protein sizes (112-172 aa) and only 40-50% identity to Cor a 1. These were labelled as ‘Cor a 1-like’ proteins, none of which has been shown to have allergenic activity.

**Figure 5.**
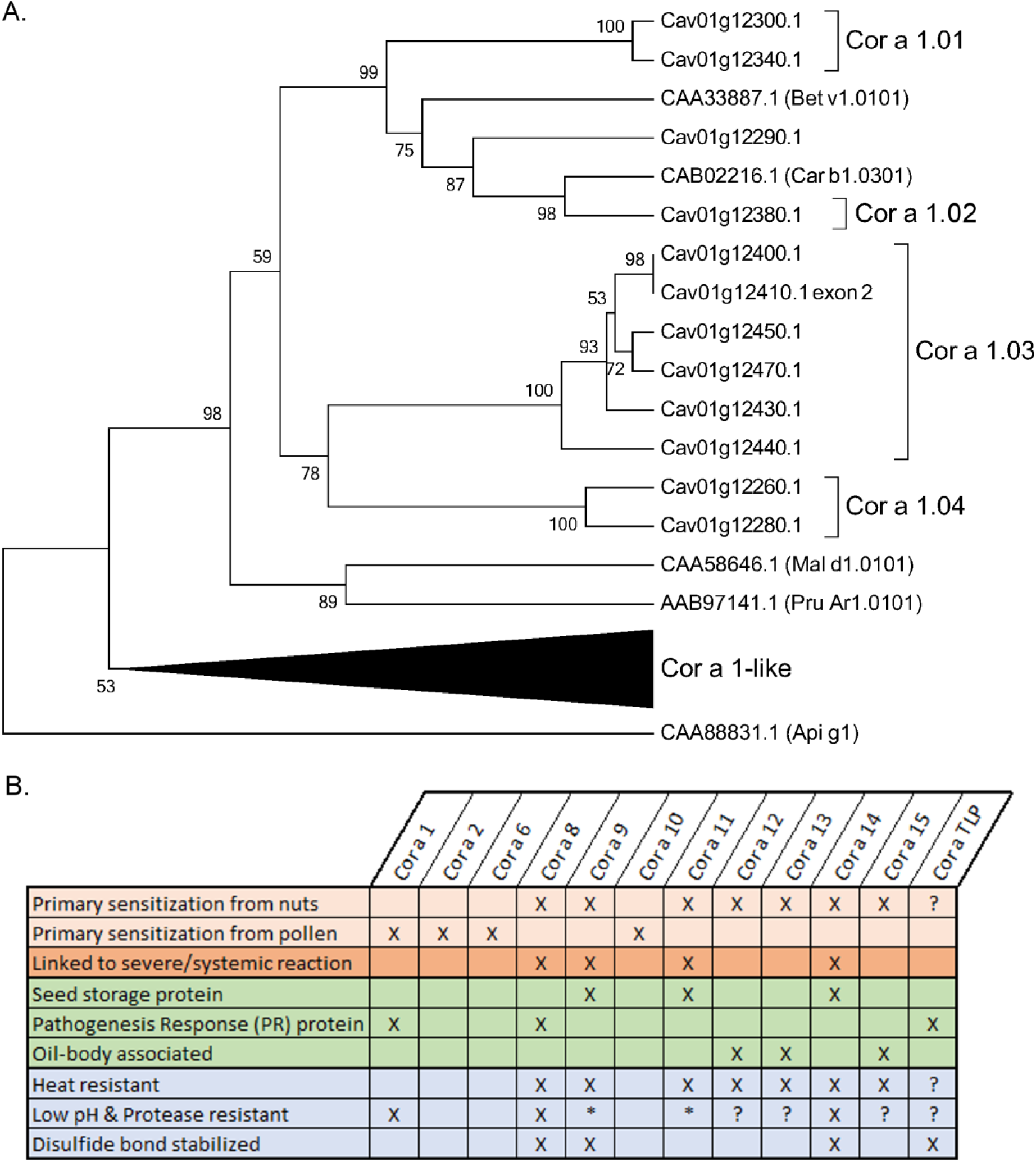
**A.** Clustering of Cor a 1 sequence homologs from the hazelnut genome, using the UPGMA method on the basis of p-distance. Node values are % of 1000 bootstrap replications. Homologs from other species of Betulaceae (Bet v 1, Car b 1) and Rosaceae (Mal d 1, Pru Ar 1) are indicated by their Genbank accession IDs. Celery allergen Api g 1 was included as an outgroup. **B.** Shared allergenic, functional and biochemical characteristics of known and suspected hazelnut allergens. *11S & 7S globulins are partially digested, but leave smaller, protease-resistant polypeptides.

From the highly conserved Cor a 1 group, two genes encoding 160 amino acid proteins are predicted to express the four isoforms of Cor a 1.01, with Cor a 1.0101 and Cor a 1.0103 most closely matching Cav01g12300, while Cor a 1.0102 and Cor a 1.0104 matched Cav01g12340. We suggest that different alleles of these two genes account for the multiple isoform variants. Cav01g12380 was the only close match to Cor a 1.02; in contrast, Cor a 1.03 had no single best match, but had >90% identity to a cluster of six genes encoding 159 amino acid proteins, making all of these potential new isoforms of this allergen. All four reported isoforms of Cor a 1.04 appear to be allelic variants of Cav01g12260, which is identical in sequence to Cor a 1.0403. However, Cav01g12280 is also >90% identical to Cor a 1.04, and so may encode additional isoforms. In addition, Cav01g12290 encodes the nearest homolog of Bet v 1.01 in the *C. avellana* genome, but falls between Cor a 1.01 & Cor a 1.02 (75% and 85% identity respectively); therefore, it expresses a putative new isoallergen, pCor a 1.05. Finally, the remaining eight Cor a 1 homologs in this locus, along with five others located on other chromosomes, formed a group of predicted proteins with more diverse sizes (112-172 aa) and only 40-50% identity to the known hazelnut allergens. These were labelled as ‘Cor a 1-like’ proteins, none of which has been shown to have allergenic activity. All putative new isoallergens and allergen-like gene models are listed in Table S14.

Genomic analysis of most of the other known hazelnut allergens was much more straightforward. Cor a 6, and Cor a 8 – 14 all matched single gene models with > 99% identity, indicating that these are the unique genes expressing these allergens. Cor a 15 is a recently reported oleosin of similar size to Cor a 12 but differing N- and C-terminal sequences (www.allergen.com). Although the protein sequence of Cor a 15 is not in the public domain at the time of writing, based on the available information Cav03g05140 is the probable gene.

Cav11g01135, which encodes Cor a 9.0101, had an adjacent 97% identical duplicate (Cav11g01145) encoding a putative new isoform; similarly, Cav06g05240 and Cav03g28030 might be isoallergens of Cor a 6 (Cav06g05250) and Cor a 14 (Cav03g29025) respectively. The profilin allergen Cor a 2 has two reported isoforms, both of which were equally close matches to Cav11g14860. However, two potential new isoallargens (Cav07g08730 and Cav11g14870) of >80% identity to Cor a 2 were also identified, and were identical in sequence to 2 profilins previously isolated from hazelnut pollen (Jimenez-Lopez et al. 2012). As with the other potential isoallergens, serological tests would be required to determine whether or not these homologs are allergenic. Remnants of a third profilin gene, which appears to have been split in half by a local chromosome rearrangement, were found in the vicinity of Cor a 2 (Cav11g14910 & Cav11g14920).

Thaumatin-like proteins (TLPs) are known to be important causes of fruit allergy, especially in the Rosaceae. Palacin et al. (Palacín et al. 2012) isolated a TLP from *C. avellana* and found cross-reactivity with sera from fruit-allergic patients, but in <10% of cases. The Genbank accession ID given for Cor a TLP in the aforementioned paper actually refers to an apple TLP; however, based on the peptide sequences also reported, we inferred that the Cor a TLP tested was encoded by Cav09g00690. This is one of 48 predicted TLPs found in the hazelnut genome; seven of these, including Cav09g00690, have >75% identity to known allergens Mal d 2 (apple) and Pru p 2 (peach). Therefore, although TLPs have not yet been demonstrated to be allergenic in hazelnut, there is a high potential for cross-reactivity between these genes and homologous fruit allergens.

In summary, we identified complete gene models encoding all known hazelnut allergens and several previously unreported putative allergenic proteins, including nine new isoforms, four new isoallergens, and suspected new cross-reactive oleosin and TLP proteins (Supplementary Table 14).

## Discussion

### Towards a reference genome for *C. avellana*

Hazelnut is typical of a number of important crop species for which, until recently, limited genome data has been available. The publicly available *C. avellana* var. ‘Jefferson’ draft genome sequence made it possible to identify the majority of gene sequences, but not their chromosomal locations. Here, we aimed to produce a reference quality genome sequence for the Turkish cultivar ‘Tombul’ in a time and cost-effective manner. This required 3 different sequencing technologies that provided data with different size ranges: 0.1-1 kbp (Illumina paired-end), 1-10 kbp (NanoPore), and 10 kbp – 10 Mbp (Dovetail). None of these methods individually or in pairs was sufficient to reconstruct the whole genome but combining all three together produced a chromosome-scale genome assembly.

The chromosomes presented here have a total length of 370 Mb, about 2.1% shorter than the estimated genome size. Each chromosome still contains several hundred small sequence gaps (Table 2), the actual size of which is not known. Also, it is likely that telomeric and centromeric repeats are more condensed in our assembly than in the physical chromosomes due to their extended repetitive structure. These two factors could explain most or all of the ‘missing’ sequence length.

Apart from these small differences, we found the chromosomes to be highly consistent with existing cytogenetic data and genetic maps, suggesting that they accurately represent the structure of the genome. Therefore, this genome assembly will be an excellent resource for accelerating breeding through novel molecular marker design and mapping candidate genes for important traits of interest, especially in ‘Tombul’, the most highly valued Turkish variety. Further high-quality assemblies from different individuals, such as that currently being constructed for ‘Jefferson’ (Snelling et al. 2018) will be invaluable for identifying the degree of variation and chromosome rearrangement within the hazelnut population.

### Comparison of the *C. avellana* genome with other horticultural crops

With the greatly reduced cost of high-throughput sequencing technologies, the genomes of a number of important nut tree species have been sequenced in recent years, such as Persian walnut, *Juglans regia* (Martínez-García et al. 2016), and pistachio, *Pistacia vera* (Zeng et al. 2019). The closest relative of hazelnut for which a complete genome is available is silver birch (*Betula pendula*), for which pseudochromosomes were generated by anchoring genome scaffolds on to a high-density genetic map (Salojärvi et al. 2017). Among these examples *C. avellana* has both the smallest genome (birch: 440 Mb, walnut & pistachio ∼600 Mb) and the lowest proportion of repetitive elements (38%; others range from 50-70% of the whole genome). These features make hazelnut an attractive model for genomic studies. While local gene replications were widespread, as in *B. pendula* there was no evidence of any recent whole-genome duplication event (Salojärvi et al. 2017). Tree species are often highly heterozygous, which means that *de novo* sequencing assemblies are often significantly longer than the expected size (Martínez-García et al. 2016; Zeng et al. 2019) due to some regions being represented twice. We observed the same effect in our assemblies that relied only on short reads; however, incorporating ∼9.3x genome coverage of long Nanopore reads reduced the assembly to the expected length, suggesting that the heterozygous regions were resolved by this method (Table 1). This gives us confidence that we can make accurate assessments of numbers of functional elements from the *C. avellana* genome.

The number of genes annotated in each of these genomes is comparable, with the 28,409 reported here being very similar to *B. pendula* (28,153) and a little less than walnut & pistachio (∼32,000 each; some of the greater number might be attributable to heterozygous duplicates). However, each of these genomes has different gene families that have undergone lineage-specific expansion, which may indicate their functional importance to their species; for example, *J*. *regia* has an unusually large complement of genes for polyphenol synthesis (Martínez-García et al. 2016). Similarly, we found 63 functional classifications (using the MapMan ontology) in which *C. avellana* had more than twice as many genes as all of the other plants examined (Supplementary Table 11). The largest groups of related functions among these classifications were components of the vesicle trafficking and RNA biosynthesis pathways; for example, HEN1, which is essential for stabilizing small regulatory RNAs (Bologna and Voinnet 2014). These observations suggest that *C. avellana* has developed diverse systems for regulating protein function at both transcriptional and post-transcriptional levels. Furthermore, 27 genes encoding triterpene synthases were identified 20 for type-II patatin-like phospholipase A2. Both of these classes of enzymes are reported to be involved in stress and defense responses in plants, the former by production of secondary metabolites (Thimmappa et al. 2014) and the latter by regulating cell death (La Camera et al. 2009). Closer examination of these gene families could reveal important aspects of the response to infection of hazelnut.

### Genomic insights into sources of powdery mildew resistance

The genome sequence enabled us to identify 11 full-length *MLO* genes in *C. avellana*, a gene family that is ubiquitous in higher plants and controls susceptibility to powdery mildew infection in diverse crop species (Acevedo-Garcia et al. 2014). In plant genomes studied to date, the *MLO* gene family varies in size from 8 (wheat) to 39 (soybean) members, and the family is divided into 6-8 clades, depending on the species included in the analysis. All *MLO* genes known to play a role in mildew susceptibility fall into Clade IV (monocots) or V (dicots). In *A. thaliana,* a loss of function mutant of *AtMLO2* confers partial resistance to mildew infection, while the triple *AtMLO2/AtMLO6/AtMLO12* mutant is completely resistant. Similarly, knockdown of *MdMLO19* in apple reduced powdery mildew infection by 75% (Pessina et al. 2016). Therefore, although sequence similarity does not guarantee conservation of function, it is likely that one or more of the hazelnut Clade V *MLO* genes could be involved in susceptibility to *E. corylacearum*. In our phylogenetic comparison (Fig. 4A), Clade VII was not clearly separated from Clade V, and Clade IV was basal to both, suggesting that these 3 may form a sub-family of *MLO* genes involved in PM susceptibility. In apple, *MdMLO19* is upregulated during PM infection, and so is *MdMLO18*, which falls into Clade VII (Pessina et al. 2014). Therefore, Cav02g29751, along with Cav07g20270, Cav07g 20290, Cav07g20321 & Cav10g10331, should be prioritized for further functional studies. In particular, it would be extremely valuable to identify natural *MLO* mutations leading to PM resistance in hazelnut germplasm, as has been documented in crops such as cucumber and apple (Berg et al. 2015; Pessina et al. 2017).

The molecular function of MLO proteins is still unclear, but in their absence mildew infection is blocked at the point of cell wall penetration (Kusch and Panstruga 2017), suggesting that mildew fungi may need to use them as a receptor to initiate cell entry. Their ubiquitous presence in plant genomes and the fact that all naturally occurring *MLO* mutants are recessive indicates that they perform a necessary function for the plant in the absence of PM. This is consistent with our observations of the genomic context of hazelnut *MLO* genes (Fig. 4B), where the Clade V/VII genes in particular seem have been selected for diversification; we hypothesize that historic PM disease pressure could have led to suppression or disruption of some *MLO* genes, while their value in the absence of disease has selected for duplication and maintenance of new gene copies. With this in mind, further study of the interaction of powdery mildew disease and the *MLO* genes in *C. avellana* would provide valuable insight into the function of this gene family in tree species.

### A catalogue of allergens suggests strategies for addressing hazelnut sensitization

In the Western world, food allergy is a widely-recognized health problem and nut allergy is one of the best studied examples, with 1-2% of the population having some kind of sensitization to hazelnut (Costa et al. 2015). A diverse group of allergenic proteins have been identified by their ability to provoke an IgE-mediated immune response in sensitized individuals (Table 4, Figure 5). We were able to identify a complete catalogue of genes for these proteins within the *C. avellana* genome; this showed that the previously reported allergen isoforms result both from multiple genes within the genome, and multiple alleles of those genes in different individuals. Based on this catalogue, we also identified likely cross-reactive allergens that have not been reported previously, which will help to guide ongoing studies aiming to treat or prevent hazelnut allergy.

The allergic response is complex and varies between individuals. For example, the allergens Cor a 1, 2, 6 & 10 are most abundant in pollen; it is thought that sensitization to these proteins primarily occurs at the mucosal membranes of the respiratory system, leading to localized symptoms. However, sensitization to consumed nuts is more likely to result in severe, systemic allergic responses; Cor a 8 and the seed storage proteins (Cor a 9, Cor a 11 and Cor a 14) have each been associated with a higher risk of severe allergy, depending on the study population (Schocker et al. 2004; Garino et al. 2010; Datema et al. 2015). Moreover, many individuals show sensitization both to pollen and nuts. This complex response makes it difficult to predict which proteins could be allergenic; however, some common features can be noted between the known hazelnut allergens (Fig. 5b). Cor a 1, Cor a 8 & TLPs are all members of ‘Pathogenesis Response’ protein families (PR-10, PR-14 and PR-5 respectively), which were first identified by their increased expression during the plant hypersensitive response to infection. They have diverse molecular functions but are relatively small proteins that resist protease degradation and are often stabilized by disulfide bonds, meaning that they are likely to be presented to the immune system with their 3-dimensional structure intact. Cor a 1, the major hazelnut pollen allergen, is known to include multiple isoallergens that cross-react with each other and those from other species (Lüttkopf et al. 2002). From the *C. avellana* genome sequence we were able to identify several new Cor a 1 variants, as well as demonstrating that they form a well conserved sub-group distinct from the other PR-10 family proteins (Fig. 5a, here described as ‘Cor a 1-like’). Given the known cross-reactivity of this class of allergens, it may be that the existence of multiple, closely related genes in the genome itself increases the risk of allergenicity. If so, the TLP family shares all of these characteristics with Cor a 1; although not conclusively proven to be allergens in hazelnut (Palacín et al. 2012), they should be regarded as high-risk. The existence of so many variants in the genome suggests that removing these allergens through breeding or genome editing would be impractical. However, Cor a 8 is a more promising target; although the nsLTPs are also a multigene family, Cor a 8 is highly diverged from the other members, so cross-reactivity is unlikely. Even so, more functional studies are needed to determine whether it is essential to the health of the tree.

The remaining nut allergens are both resistant to the heat used in cooking and the acidity and protease activity of the gastrointestinal tract. The seed storage proteins are highly abundant, making up >50% of all protein in nuts (Beyer et al. 2002). They are also found in condensed ‘protein bodies’ within the plant cell, increasing the probability that at least some of these proteins will be presented to the immune system with their conformational epitopes intact. Similarly, the oleosins are tightly associated with intracellular oil droplets, which may help to protect them from degradation (Akkerdaas et al. 2006). They were only recognized as allergens relatively recently, because methods used to produce protein extracts from nuts often eliminate oil droplets; however, Cor a 12-sensitivity was observed consistently in 10-25% of hazelnut allergic patients across Europe (Datema et al. 2015).

This illustrates that, while empirical testing of allergic response is essential to characterize allergens, there is always a risk of overlooking some important factors. In contrast the genomic survey presented here can give confidence that all relevant proteins have been identified and provide a foundation for further studies of allergenicity.

### Conclusions

We present here a chromosome-level reference genome assembly and annotation for European hazelnut, *C. avellana* cv. Tombul. Using a combination of short-read, long-read and proximity ligation sequencing we produced a genome of similar quality to those obtained by anchoring contigs to high-density genetic maps, making this to our knowledge the most complete tree nut genome published to date. The genes and functional elements identified here provide a foundation for ensuring the sustainability of future hazelnut production, for example by identifying targets for breeding or gene knockout that could confer resistance to powdery mildew disease and decrease the risk of hazelnut allergy.

## Methods

### Plant materials and DNA purification

2-year old saplings of *C. avellana* L. var. Tombul were obtained from commercial nurseries in Turkey and cultivated on the Sabanci University campus. Isolation of high-quality gDNA proved to be difficult, due to the abundance of polysaccharides and other compounds in hazel tissues that are not easily separated from DNA by standard techniques. Therefore, we adopted an isolation method previously developed for *Betula nana* (Wang et al. 2013), with some modifications: best results were obtained by isolating DNA from leaf buds, the incubation time with 2x CTAB buffer was shortened to 1 hr at 65°C, and RNAse A was applied in this step rather than as a separate incubation. Purified DNA was additionally bound to a silica membrane (NucleoSpin Plant II Kit, Macherey-Nagel, Düren, Germany) and washed to remove low molecular weight DNA fragments, before being eluted in 60 µl of TE buffer. Final DNA concentration was measured using a dsDNA-specific fluorescent dye (Quant-iT HS dsDNA Assay Kit, ThermoFisher, Waltham, MA, USA).

### Next-Generation Sequencing & d*e novo* genome sequence assembly

Illumina library preparation and sequencing were carried out by Macrogen (Seoul, S Korea). 2 Paired-end shotgun sequencing libraries were produced using TruSeq Library Preparation kits, size selected to have an average insert of 700-800 bp, and sequenced on a single lane of a HiSeq 4000 instrument (Illumina, San Diego, CA, USA). Single-molecule sequencing was carried out in-house; whole genomic DNA was physically disrupted into ∼8kb fragments using a Covaris g-TUBE (Covaris, Woburn, MA, USA) and then prepared for NanoPore sequencing on the MinION platform using the Ligation Sequencing Kit 1D, according to the manufacturer’s protocols (Oxford NanoPore Technologies, Oxford, UK). Data was obtained from a 48 hr run on a single R9.4 flowcell. Proximity ligation sequencing was carried out by Dovetail Genomics (Santa Cruz, CA) using their proprietary Chicago & HiC protocols.

The quality of high-throughput sequencing data was assessed using FastQC (Andrews and Babraham Bioinformatics 2010). *De novo* sequence assembly was carried out on the High Performance Computing cluster (HPC) at Sabanci University. For Illumina-only assembly, raw sequence reads were processed with Trimmomatic (Bolger et al. 2014) to remove TruSeq adapters, trim bases with quality score <5 from both ends of the reads, and use a sliding window of 4 bases to cut the reads when average sequence quality across the window dropped below 20. The trimmed sequences were then assembled using ABySS 1.9 (Simpson et al. 2009) with a range of k-mer size values; k=80 was found to empirically to give the most contiguous assembly, which was improved further by enabling scaffolding across large bubbles in the k-mer graph (POPBUBBLES_OPTIONS=--scaffold b=5000). MinION sequencing adapters were trimmed from the first and last 50 nt of NanoPore reads using bbduk from the BBtools suite (Bushnell 2016) with the options k=19, editdistance=3. Hybrid genome assembly of the Illumina and trimmed NanoPore reads was carried out using MaSuRCA 3.2 (Zimin et al. 2013), with the average Illumina insert size specified as 790±80 nt. This assembly took approximately 9 days running on 36 CPUs in parallel. Proximity ligation data was integrated with both the Illumina-only and the Hybrid genome assembly using Dovetail Genomics’ HiRise assembly pipeline.

### Assessment of assembly completeness and genome sequence comparisons

The completeness and accuracy of genome assemblies was assessed using BUSCO v3 (Waterhouse et al. 2018) using default settings for the provided virtual machine, with the reference database for single copy genes found in land plants (embryophyta) from OrthoDB v9.1(Kriventseva et al. 2015). The draft *C. avellana* cv. ‘Jefferson’ genome, CDS and annotations were retrieved from the original producers’ web portal (http://hazelnut.data.mocklerlab.org/). Tree genome assemblies used for inter-species comparison were as follows (with GenBank Assembly ID): *Betula nana* (GCA_000327005.1), *Betula pendula* (GCA_900184695.1), *Populus trichocarpa* (GCA_000002775.3), *Prunus persica* (GCA_000346465.2), *Juglans nigra* (GCA_003123865.1) and *Juglans regia* (GCF_001411555.1).

Routine sequence similarity searches for single or moderate numbers of sequence elements in the genome assemblies were carried out using standalone BLAST+ 2.2.30 (Camacho et al. 2009). Whole genome searches and comparisons were realised with bwa 0.7.12 (Li and Durbin 2010) and resulting alignment files were processed and analysed using SAMtools 1.8 (Li et al. 2009) and bcftools 1.8 (Danecek et al. 2011).

### Detection and masking of repetitive elements

Repetitive elements were identified using RepeatMasker 4.0.7 on ‘normal’ sensitivity with default scoring matrices and a custom repeat database produced by combining novel repeats detected in the Tombul genome (Supplementary Information) with all eudicot repetitive elements recorded in RepBase Update 22.08 (5,913 elements) and mipsREdat 9.3 (26,123 elements) (Jurka et al. 2005; Smit et al.; Nussbaumer et al. 2012). Detected repeats were masked with runs of ‘N’. For subsequent identification of protein-coding genes and other functional elements, the `–nolow` option was used to leave simple repeats and low-complexity regions unmasked.

### Prediction of hazelnut functional RNAs and miRNA targets

Discovery of tRNA genes was carried using tRNAscan-SE 2.0.0(Chan and Lowe 2019). To detect rRNA, masked chromosome sequences were searched using BLASTN (e-value cut-off 1e-30) with the following coding sequences: previously published 5S rRNA of *C. avellana* (Genbank HF542974.1(Falistocco and Marconi 2013)); and complete tree 45S rRNA sequences retrieved from Genbank.

Conserved miRNA genes were identified using all reported plant miRNAs from miRBase v22 (Kozomara et al. 2019) with SUmirFind and SUmirFold (Lucas and Budak 2012), using a mismatch cutoff ≤ 3 for initial miRNA homolog detection, followed by predicted pre-miRNA secondary structure prediction and selection of strong miRNA candidates based on established structural criteria (Lucas and Budak 2012). For inter-species comparison, the same methods were used to identify miRNA genes in other plant genomes.

Expression of putative miRNAs was confirmed by using all non-redundant pre-miRNA sequences to search assembled Tombul transcripts using BLASTN, retaining hits with >78% identity, >80% coverage as expressed pre-miRNAs. Experimentally validated targets of specific miRNA families found in *C. avellana* were retrieved from miRBase. Predicted targets of putative miRNAs were found in the transcripts using psRNATarget (Dai et al. 2018). Predicted targets were annotated with Gene Ontology terms using Blast2GO software (Conesa and Götz 2008).

### Protein-coding Gene Modelling and Annotation

Prediction of gene models was carried out by Augustus, an *ab initio* gene predictor based on Hidden Markov Models (Stanke and Waack 2003). Augustus was run on the masked Tombul genome using parameters optimised for *Arabidopsis thaliana*. High-confidence genes were then identified by aligning gene models to the Tombul transcriptome using BLASTN, and retaining hits with ≥90% sequence identity and ≥25% query coverage. Additional gene modelling in genome regions with significant homology to orthologs from other plants (>50% identity over >35 amino acids) was carried out using FGENESH (Solovyev et al. 2006) with training parameters from *Betula nana*. Functional annotation of the gene models was carried out using Mercator, Mercator4, and TRAPID with default parameters, and additional GO term plots were produced using REVIGO (Schwacke et al. 2019; Van Bel et al. 2013; Supek et al. 2011).

### Characterization of MLO and allergen gene families

All gene models annotated with the InterPro domains IPR004326, ‘Mlo-related protein,’ IPR000916, ‘Bet v I allergen’, and other allergen-related domains were inspected individually. *A. thaliana* MLO protein sequences were obtained from the Araport11 genome annotation (Cheng et al. 2017), while those for *Malus domestica* came from the Genome Database for Rosaceae (Jung et al. 2019). Reference allergen sequences were retrieved from Genbank, using the accessions listed at www.allergen.org. *MLO* and allergen gene models were verified and re-annotated as described in Supplementary Information. Transmembrane structure of predicted MLO protein sequences was evaluated using TMHMM 2.0 with default parameters (Krogh et al. 2001). Incomplete gene fragments and unexpressed homologs were detected by searching the pseudochromosomes using tblastn, with the complete protein sequences as the query. MEGA 6.0 (Tamura et al. 2013) was used for multiple sequence alignments (using the Muscle algorithm) and for phylogenetic clustering.

### Data Access

All raw and processed sequencing data generated in this study have been submitted to the European Nucleotide Archive (ENA), under accession number PRJEB31933 (https://www.ebi.ac.uk/ena/data/view/PRJEB31933).

## Acknowledgments

This work was supported financially by the Scientific and Technological Research Council of Turkey (TÜBITAK grant no. 215O446 to SJL) and by the Newton Fund Institutional Links programme (Grant no. 216394498 to RJAB). This study utilized the Sabanci HPC Cluster for computing support. The authors would like to thank Nihal Öztolan Erol and Andrew J. Helmstetter for helpful discussions.

## Author Contributions

SJL conceived the study, developed analysis pipelines, analyzed and interpreted data and wrote the manuscript. KK & BA acquired, analyzed and interpreted data, prepared figures, and drafted sections the manuscript. RJAB contributed substantively to study design and revised the manuscript. IB analyzed and interpreted additional data. All authors read and approved the final manuscript.

